# Perturbations in podocyte transcriptome and biological pathways induced by FSGS associated circulating factors

**DOI:** 10.1101/2022.09.08.507168

**Authors:** Priyanka Rashmi, Tara K. Sigdel, Dmitry Rychkov, Izabella Damm, Andrea Alice Da Silva, Flavio Vincenti, Andre L. Lourenco, Charles S. Craik, Jochen Reiser, Minnie M Sarwal

**Author notes:** Joint first authors (Priyanka Rashmi, Tara K. Sigdel). Corresponding author: Minnie Sarwal, MD, PhD, MRCP, FRCP, Professor in Residence, Surgery, UCSF, Co-Director, Kidney Pancreas Transplant Program, UCSF. Author Contributions: Manuscript was drafted by PR, TS, DR, MS; MS, TS, ID, PR designed the study; FV, MS, CC, AL contributed to sample collection and clinical data capture; TS, PR, ID, DR, AS, AL generated and analyzed the data. All authors (PR, TS, DR, ID, AS, FV, AL, CC, JR, MS) listed contributed to the data analysis and/or interpretation, reviewed and approved the manuscript including the final version as submitted.

## Abstract

Focal segmental glomerulosclerosis (FSGS) is frequently associated with heavy proteinuria and progressive renal failure requiring dialysis or kidney transplantation. However, primary FSGS also has 40-80% risk of recurrence of disease in the transplanted kidney (rFSGS). Multiple circulating factors have been proposed to contribute to the pathogenesis of primary and rFSGS. However, neither the factors nor the downstream effectors specific to individual factors have been identified. The tumor necrosis factor, TNF pathway activation by one or more circulating factors present in the sera of patients with FSGS has been supported by multiple studies. The proposed circulating factors include soluble urokinase-type plasminogen activator receptor (suPAR) and patient derived CD40 autoantibody (CD40autoAb) in the development and recurrence of FSGS. In a human *in vitro* model, using two novel human antibodies-anti uPAR (2G10) and anti CD40 antibody, we show that the podocyte injury caused by sera from FSGS patients is at least in part mediated by CD40 and suPAR. Additionally, we employ gene expression studies to compare the molecules and pathways activated in response to CD40 autoantibody from rFSGS patients (rFSGS/CD40autoAb) and suPAR, and delineate the unique pathways associated with FSGS injury and transcriptional podocyte alterations with targeted blockade of suPAR and CD40 pathways.

**Clinical Impact:** Focal Segmental Glomerulosclerosis remains a disease without specific therapy for primary disease and high rate of recurrence after kidney transplantation. Circulating factors are implicated in the pathogenesis of FSGS but targeting them for therapy has remained elusive. We propose two potential therapeutic molecules for rFSGS treatment-a human anti-uPAR antibody (2G10) and a humanized anti-CD40 blocking antibody (Bristol Meyer Squibb, 986090) that reverse podocyte injury associated with FSGS in cultured podocytes and can be further tested in pre-clinical and clinical models. Furthermore, we use microarray profiling to identify transcriptional pathways specific to podocyte injury from patient-derived CD40 autoantibodies (rFSGS/CD40autoAb) and suPAR and selective blockade of these pathways to abrogate podocyte injury.

## Introduction

Focal segmental glomerulosclerosis (FSGS) is the most common primary glomerular disorder causing end-stage renal disease in the United States [1]. It is frequently associated with severe proteinuria (>3.5 g/day in adults and >1 g/day in children), scarring of the glomerulus and loss of terminally differentiated podocytes. The etiology of FSGS is diverse including genetic mutations in podocyte genes, adverse drug interactions, toxic insults and viral infections. However, in the majority of FSGS cases, the etiology remains unknown (idiopathic). Whatever the cause, the injury is primarily directed at the podocytes [2]. Podocytes are highly differentiated cells of the renal glomerulus consisting of a cell body, major processes and foot processes (FPs). FPs form a characteristic interdigitating pattern with FPs of neighboring podocytes, leaving in between the filtration slits that are bridged by the glomerular slit diaphragm [3]. The slit diaphragm and the apical and basal membrane domains of podocytes are connected to each other by an actin-based cytoskeleton, which is critical for the maintenance of the glomerular filtration barrier. It is reported that idiopathic FSGS results from disorganization of the podocyte cytoskeleton and slit diaphragm with consequent foot process effacement, podocyte hypertrophy, detachment from the GBM and loss with migration into the Bowman space [4]. Idiopathic FSGS recurs after transplant in about 40% of adult and pediatric patients, in some cases within a few hours or days after kidney transplantation [5]. These clinical observations have given rise to the idea that FSGS is associated with circulating factors generated after cellular or humoral immune reactions.

Some of the proposed circulating factors that may increase glomerular permeability are soluble urokinase-like plasminogen activator receptor (suPAR), cardiotrophin-like-cytokine-1 (CLC-1), vascular permeability factor (VPF) and hemopexin [6]. We recently described a panel of autoantibodies in sera from FSGS patients to predict the risk of recurrence after transplant [7]. Furthermore, we showed interaction between two proposed circulating factors in augmenting podocyte injury *in vitro* and *in vivo*. Administration of suPAR and anti CD40 antibodies isolated from rFSGS patients (rFSGS/CD40autoAb) caused an increase in proteinuria in mice and enhanced podocyte injury in an *in vitro* cell culture model of immortalized human podocytes [7, 8]. Our study suggested that anti-uPAR and anti-CD40 therapies may be effective in slowing down or reversing the renal injury leading to recurrence after transplant.

CD40 is a costimulatory molecule of the tumor necrosis factor (TNF) receptor (TNFR) superfamily. CD40 and its ligand (CD40L/CD154) play a fundamental role in both humoral and cell-mediated immunity [9]. CD40 is constitutively expressed on professional antigen-presenting cells (B cells, dendritic cells, and macrophages) and can be induced on several non-hematopoietic cells including the parenchymal cells of the kidney [10]. CD40L is expressed mainly by activated T cells and platelets but can also be secreted into the blood by platelets as a soluble protein [11]. A role for CD40/CD40L signaling in the development of proteinuria in various disease settings has been demonstrated. sCD40L has been shown to directly act on podocytes and cause loss of nephrin in culture. Additionally, sCD40L increases albumin permeability in isolated rat glomeruli that can be inhibited by pre-treatment with inhibitors of CD40/CD40L interaction [12]. Recently, the efficacy of an anti-mouse CD40 antagonist antibody in reversal of proteinuria in two Systemic Lupus Erythematosus (SLE)-prone mouse strains was demonstrated [10].

The urokinase plasminogen activator receptor (uPAR) is a membrane-bound receptor for urokinase plasminogen activator (uPA) which regulates plasminogen-mediated extracellular proteolysis. uPAR has three domains and suPAR represents a soluble form of uPAR [13]. uPAR is expressed on several cell types such as active leukocytes, endothelial cells and podocytes. The role of suPAR in the development of FSGS has been extensively studied. In cultured podocytes, membrane-bound uPAR binds and activates αvβ3 integrin and small GTPases Rac1 and Cdc42 promoting motility [14]. Similarly, suPAR binds and activates β3 integrin in podocytes and high doses of suPAR in mice cause proteinuria [15]. This podocytopathic effect of suPAR is dependent upon its ability to bind and activate β3 integrin as no proteinuria and foot process effacement was observed in mice expressing suPAR that is incapable of β3 integrin binding (*sPlaur_E134A_*) [15].

In this study we have tested and confirmed the ability of two novel humanized antibodies-anti uPAR (2G10) and anti CD40 (kindly provided by Bristol Meyer Squibb, 986090), to reverse podocyte injury in an *in vitro* model of cultured human podocytes. 2G10, was isolated from a fragment-antigen binding (Fab) phage display library using soluble, recombinant human uPAR as the antigen to biopan the library [16]. This antibody competes with uPA for uPAR binding to inhibit cell invasion *in vitro* and in a preclinical model of aggressive breast cancer [17, 18]. BMS-986090 is a dimeric anti-human CD40 VH antagonist domain antibody formatted with a human IgG4 Fc tail (dAb-huIgG4), and an antagonist for CD40-CD40L mediated signaling. We showed that anti-uPAR and anti-CD40 antibodies are both efficacious in inhibiting podocyte injury caused by sera from different rFSGS patients, highlighting their role in FSGS pathogenesis. In addition, these results also suggest that multiple factors might contribute to and/or synergize in the development and/or maintenance of podocyte injury in primary FSGS. A combination of one or more circulating factors in FSGS has been proposed as triggers for the early activation of the TNF pathway resulting in podocyte injury [19]. Elevated serum TNF levels have been described in patients with nephrotic syndrome [20] and a beneficial role for

TNF suppression has been observed in a subset of pediatric FSGS patients [21]. While glomerular TNF but not serum TNF is associated with loss of eGFR, activation of the TNF pathway in podocytes treated with sera from FSGS patients has been demonstrated repeatedly [22, 23]. We have performed a global transcriptomic profiling of podocytes treated with two proposed circulating factors in FSGS-rFSGS/CD40autoAb, suPAR and combination of rFSGS/CD40autoAb with suPAR and compared them to TNF treated podocytes to identify common and unique pathways for these different podocytopathic factors, and their potential contribution to FSGS progression and recurrence.

## Materials and Methods

### Collection of patient sera

Sera from FSGS patients, healthy individuals and patients with end-stage renal disease due to non-FSGS related causes (non-FSGS) were collected at the University of California San Francisco (UCSF) under institutional board-approved protocols. FSGS patients were further characterized as primary FSGS, based on the exclusion of causes of secondary FSGS, and biopsy confirmed diagnosis of FSGS recurrence after transplantation, together with clinical FSGS recurrence with proteinuria, edema and hypoalbuminemia [1]. The demographic information for all the patients is provided in **Table S1**.

### Measurement of suPAR, anti CD40 antibody and total IgG content in serum

Anti-CD40 antibody in sera was measured using a previously optimized Meso Scale Discovery platform [7]. Briefly, patient serum was added to a 96-well MULTI-ARRAY plate (Meso Scale Discovery) coated with CD40 protein (Abcam) and allowed to bind at room temp for 2hr. After washing, SULFO-tag anti-human secondary antibody (MSD) was added, and the plate was read on a MESO Quickplex imager (Meso Scale Discovery, Rockville, MD). suPAR levels were measured using a commercial quantikine ELISA kit (R&D Systems). Total IgG in human sera was measured using a commercial human IgG ELISA kit (Fisher Scientific) according to manufacturer’s protocol.

### Cell culture and treatment with anti uPAR or anti CD40 antibodies

Immortalized human podocytes have been previously described [24]. Briefly, primary human podocytes were immortalized by transfection with the temperature sensitive SV40 T gene. These cells proliferate at the “permissive” temperature (33°C) and are considered undifferentiated. After transferring to the “nonpermissive” temperature (37°C), they enter growth arrest and by day 10-14 express markers of differentiated podocytes *in vivo*, such as nephrin, podocin, CD2 associated protein (CD2AP), synaptopodin, and known molecules of the slit diaphragm ZO-1, α, β, and γ-catenin, and P-cadherin. The podocytes were cultured in RPMI medium supplemented with insulin, transferrin, selenium, sodium pyruvate (ITS-A, Gibco #513000), 10% FBS and penicillin/streptomycin. After differentiation, cells were either left untreated or treated with 4% sera from patients for 24 hours. For rescue of stress fibers, podocytes were treated with patient sera in the presence of a control antibody (human IgG, Invitrogen #02-7102) or anti uPAR antibody (2G10) or anti CD40 antibody (BMS, 986090) at 1μg/mL for 24h. The 2G10 antibody was prepared as previously described [16]. Following treatment, podocytes were fixed in 4% paraformaldehyde (PFA)/Sucrose, permeabilized with 0.3% Triton X-100 and stained with rhodamine-conjugated phalloidin to visualize stress fibers. Cells were imaged by Lecia SP5 confocal microscopy and number of cells with intact stress fibers was counted.

### Podocyte treatment with proposed circulating factors

Immortalized podocytes were cultured and differentiated as described above. Primary podocytes were isolated from healthy renal nephrectomy tissue as described previously [24]. CD40Ab was purified from pooled plasma of rFSGS patients (rFSGS/CD40autoAb). Differentiated podocytes were treated with patient purified CD40Ab alone (10μg/mL) or together with suPAR (1μg/mL) or a commercially available mouse monoclonal CD40Ab (R&D mAb, 5ug/mL) for 24h. Podocytes were also treated with suPAR alone (1μg/mL), TNFα (0.1μg/ml) for 24h. **Figure 1** outlines the study design.

**Figure 1.**
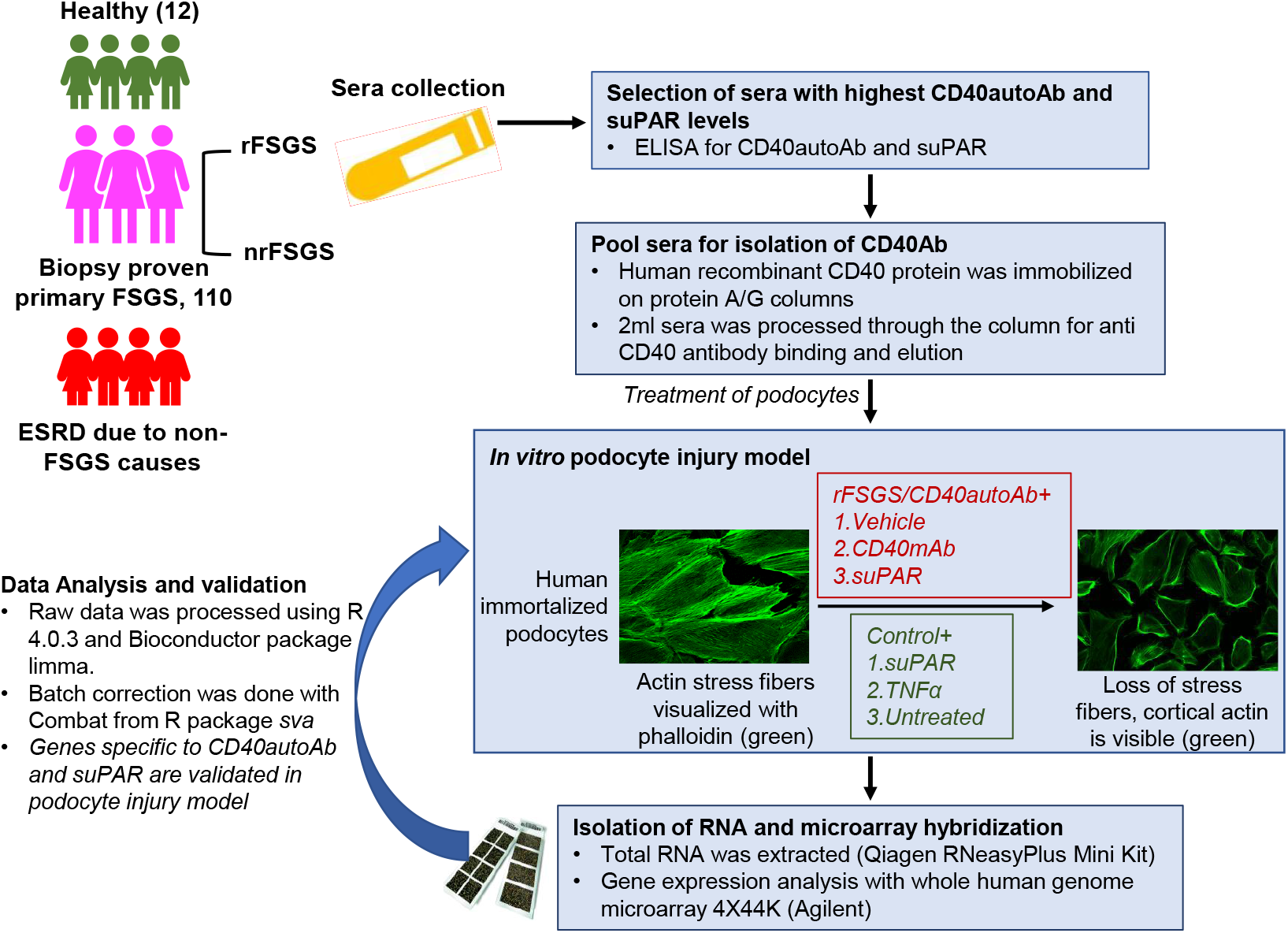
Study design for the transcriptomic profiling of podocytes. Serum/plasma was collected from end-stage renal disease (ESRD) patients with biopsy proven primary FSGS or ESRD due to non-FSGS causes. Primary FSGS group included patients that underwent biopsy proven recurrence of FSGS (rFSGS) and patients that did not recur after renal transplantation (nrFSGS). Sera from healthy individuals was collected as control. Levels of CD40atuoAb and suPAR were measured with ELISA and rFSGS patients with highest CD40autoAb levels were selected for further experiments. For the transcriptomic profiling, CD40autoAb was purified from pooled sera of rFSGS patients with highest autoantibody levels (rFSGS/CD40autoAb). Differentiated podocytes in culture were treated with rFSGS/CD40autoAb (10ug/mL) in the presence of suPAR (1ug/mL), or commercial mouse monoclonal anti CD40 antibody (CD40mAb, R&D Tech, 5ug/ml) or suPAR alone, TNF alone or left untreated. RNA was extracted according to the manufacturer’s protocol and subjected to Agilent 44k microarray. For validation, differentiated human podocytes in culture were treated with sera from patients or healthy control and loss of stress fibers was rescued with novel blocking antibodies against uPAR and CD40.

### RNA isolation and microarray hybridization

RNA was extracted from podocytes using the RNeasyPlus Mini Kit (Qiagen) following manufacturer’s instructions. RNA integrity was assessed spectrophotometrically using NanoDrop (Thermo Scientific) and ensuring that the 260/280 ratio was approximately 2.0. A whole Human Genome Microarray 4X44K (Agilent Technologies) was used for gene expression analysis. Images were scanned with a dual-laser microarray scanner (Agilent C, Agilent Technologies) and data was extracted using Feature Extraction (FE) software (Agilent Technologies).

### Data Processing

The raw data were processed using R language version 4.0.3 [25] and the Bioconductor [26] package limma [27]. Processing steps included background correction, log2-transformation, and Loess normalization. The cell culture source and day of microarray hybridization were recognized as a source of non-biological data variability using principal component analysis (PCA). The combat approach from the R package *sva* [28] was used to eliminate the technical factors.

### Differential gene expression analysis

The differential gene expression analysis was performed using the R package *limma* [27]. The cell culture source and microarray hybridization date were used as covariates in the linear model. Differentially expressed genes were identified using a p-value less than 0.05 adjusted with the Benjamini-Hochberg multiple testing correction approach and > 1.2-fold change.

### Pathway Analysis

The gene sets were analyzed for enrichment in Gene Ontology biological pathways using the R package *clusterProfiler* [29]. Genes with more than 1.2 log fold change were included in the analysis and enriched pathways with q-value < 0.05 were accepted as significant.

## Results

### suPAR and anti CD40 antibody levels are higher in pre transplant sera from rFSGS patients

suPAR, anti CD40 antibody and total human IgG concentration in sera of each patient were measured as described and the values are provided in **Table 1**. Both suPAR and anti CD40Ab levels were at least two-fold higher in each patient as compared to that reported in healthy individuals ([30], Rashmi et al, unpublished results).

**Table 1.** CD40 auto-antibody (CD40autoAb) and suPAR levels in sera from FSGS patients with biopsy-proven recurrence after kidney transplant.

|  | Patient 1 | Patient 2 | Patient 3 | Mean level reported in healthy individuals |
| --- | --- | --- | --- | --- |
| suPAR (ng/mL) | 5.17 | 8.934 | 6.806 | 2.1 |
| CD40 (ng/ul) | 563.03 | 1041.89 | 980.55 | 214.1 |
| Total IgG (ng/ul) | 5253.69 | 6208.20 | 11626.74 | 8324.01 |
| CD40 /IgG | 0.11 | 0.17 | 0.08 | 0.02 |

### Co-treatment with anti uPAR antibody or anti CD40 antibody inhibits podocyte injury caused by sera from rFSGS patients

Differentiated podocytes were treated with sera (4% final concentration in media) from patients with rFSGS after kidney transplantation (n=3) and controls with non-recurrent FSGS (nrFSGS, n=1) and end stage renal disease due causes other than FSGS (non-FSGS, n=1) for 24 hours. The podocyte actin cytoskeleton was visualized by labeling with rhodamine-conjugated phalloidin as described (methods). Cells were imaged by confocal microscopy and the number of cells with intact stress fibers were counted. As shown in **Figure S1**, sera from rFSGS but not controls (nrFSGS and non-FSGS patients), caused a 50% reduction in the percentage of cells positive for stress fiber. Next, differentiated podocytes were treated as above with sera from rFSGS patients along with a human IgG for control or anti-CD40 antibody (Bristol Meyer Squibb, 986090) or anti-uPAR antibody (2G10) and stress fiber positive cells were counted as before. As shown in **Figure 2**, 2G10 was able to rescue the loss of stress fibers in all rFSGS sera treated podocytes, restoring podocyte morphology to that seen in untreated/control sera treated cells. Similarly, the number of stress fiber positive cells upon co-treatment with anti-CD40 antibody and sera from rFSGS patients, was higher than podocyte treatment with patient serum alone (81.1±10.9 vs 53.4±12.2). Control human IgG had no effect on the number of stress fiber positive cells (**Figure S1**). These results support a role for CD40 and uPAR signaling in the podocytopathy of rFSGS.

**Figure 2.**
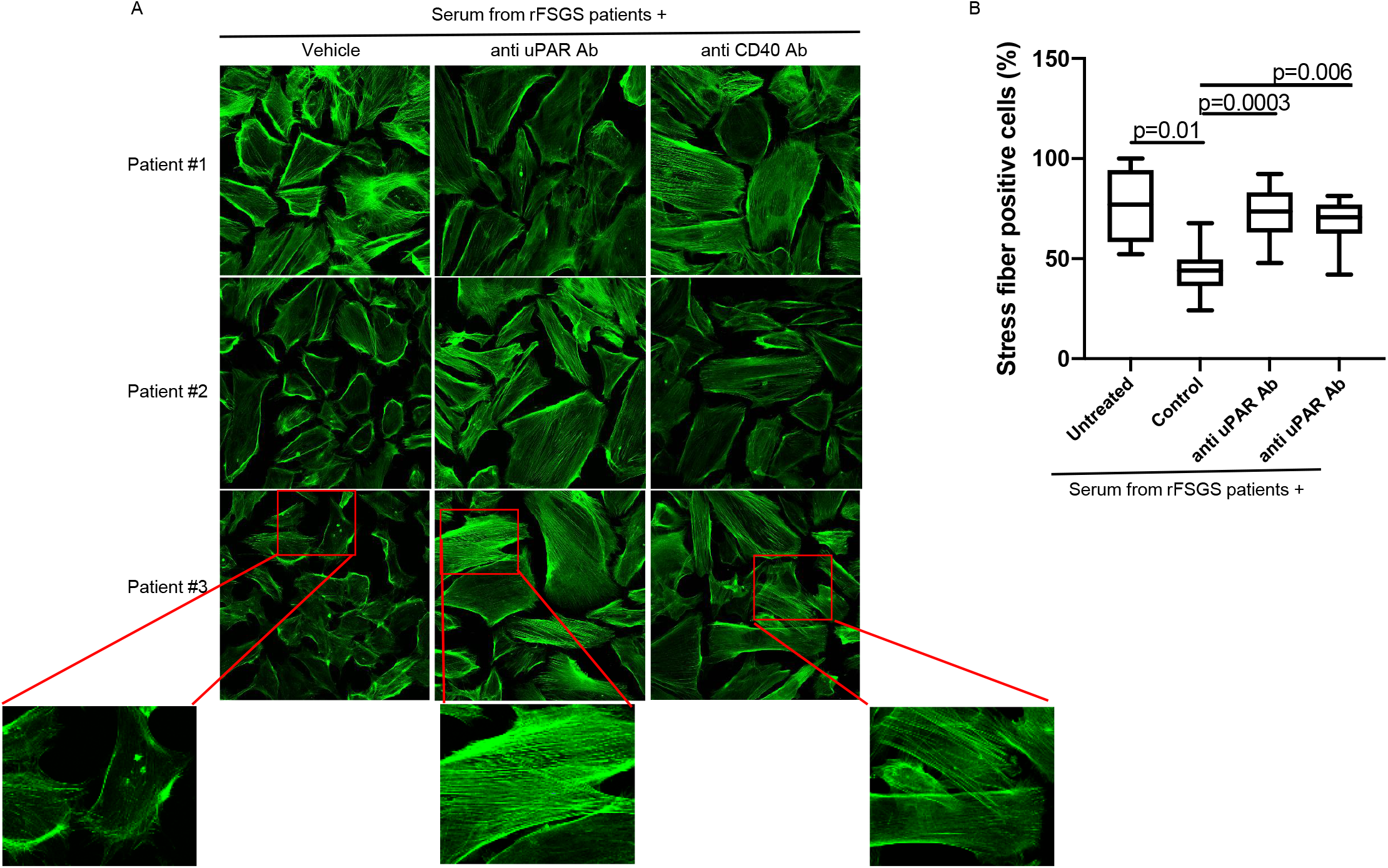
Blocking anti CD40 and anti uPAR antibodies rescue the actin depolarization in podocytes induced by sera from FSGS patients with recurrence of FSGS after renal transplant (rFSGS). A) Representative images displaying stress fibers in cultured human podocytes stained with rhodamine conjugated phalloidin (green) after treatment with pretransplant sera from three rFSGS patients. Podocyte were cultured in the presence of sera (4% final concentration) with or without pre-treatment with anti uPAR or anti CD40 blocking antibody. B) Quantification of stress fiber positive podocytes (minimum 100 cells per condition were counted). Treatment with sera from rFSGS patients caused significant depolarization of stress fibers as determined by number of stress fiber positive cells (30%, 59% and 49% reduction with respected to untreated podocytes respectively). Pretreatment of podocytes with novel blocking anti CD40 antibody or anti uPAR antibody (1ug/mL) before addition of patient sera rescued stress fibers.

### Identification of genes and pathways dysregulated in response to specific circulating factors

Recently it has been proposed that regardless of the nature of one or more circulating factors present in the serum of patients with FSGS, the renal intrinsic activation of the TNF pathway contributes to disease pathogenesis and/or progression of FSGS [22]. We have previously shown that suPAR and anti-CD40 autoantibody isolated from FSGS patients (rFSGS/CD40autoAb) alone or in combination can cause podocyte injury *in vitro* as well as in mice. However, the injury was more pronounced in mice when suPAR and rFSGS/CD40autoAb were used in combination [7]. Therefore, we sought to deconvolute the pathways and markers associated with each of the proposed circulating factors by microarray analysis. **Supplementary Figure S2** shows that the podocyte injury response to rFSGS/CD40autoAb is CD40 signaling specific as it can be reversed by a commercial monoclonal anti-CD40 antibody (CD40mAb) that blocks CD40 signaling. We found differential expression of 645 genes in response to treatment with patient derived rFSGS/CD40autoAb. Upregulated genes such as *ISG15, IFI27* and *LCN2* (**Figure 3A**) are involved in the regulation several pathways including type I interferon signaling pathway, cytokine-mediated signaling pathway as well as endothelial and epithelial cell migration (**Figure 3B**). suPAR treatment alone resulted in the differential expression of 1,737 genes primarily involved in the negative regulation of megakaryocyte development, cell surface receptor signaling and DNA replication dependent nucleosome assembly among others (**Figure 3A and 3B**). This is consistent with a major role for suPAR in the platelet biology.

**Figure 3.**
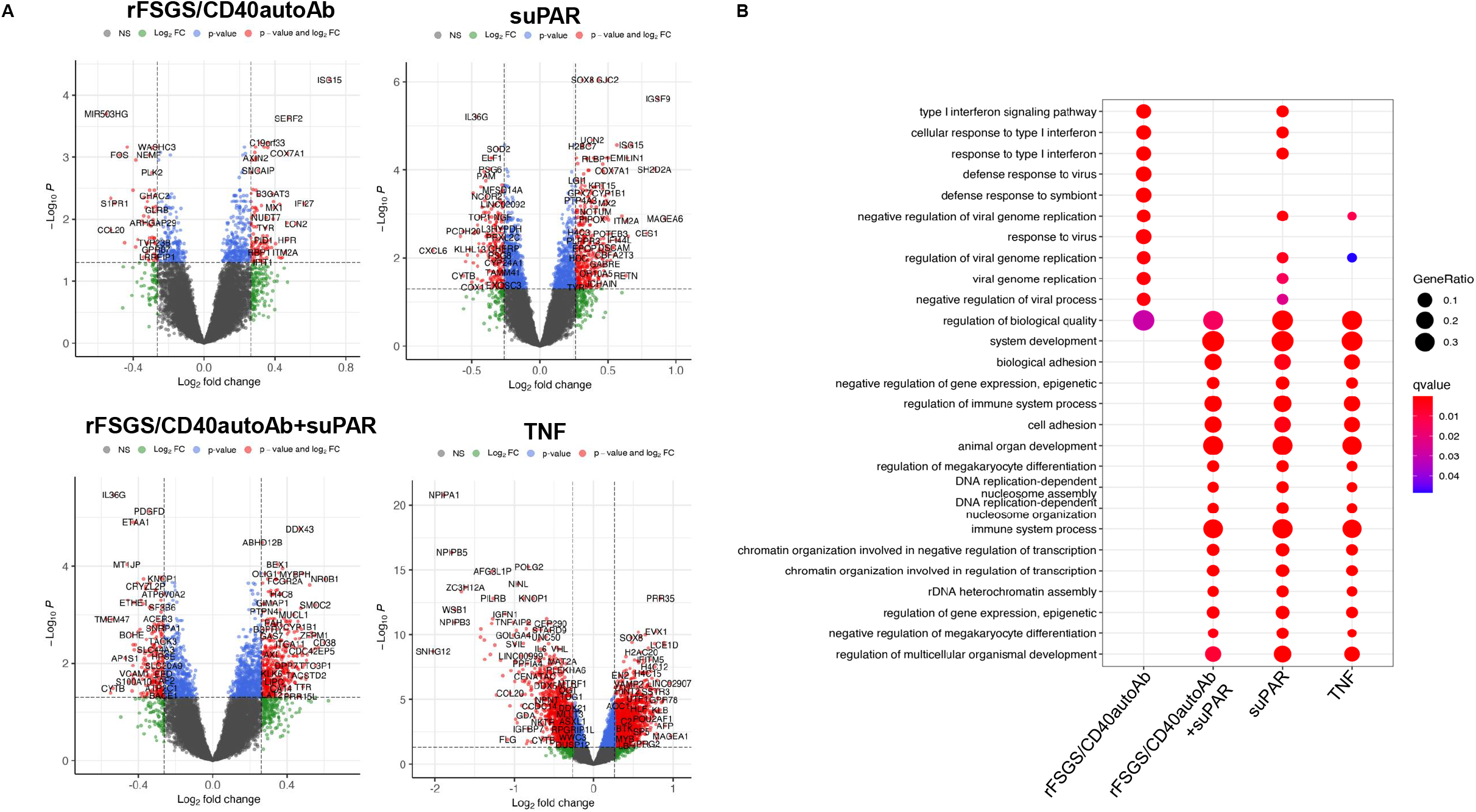
Identification of molecular pathways activated in podocytes exposed to various stimuli. Microarray was performed on RNA isolated from cultured human podocytes exposed to CD40autoAb derived from FSGS patients with recurrence of FSGS after kidney transplant (rFSGS/CD40autoAb), suPAR, a combination of suPAR and rFSGS/CD40autoAb or TNF, as described in materials and methods, was conducted. Raw data was processed using R language version 4.0.3 to obtain gene expression profiles after each treatment followed by differential gene expression and gene expression KEGG pathway analysis. A) For each treatment volcano plot of significantly altered genes with greater than 1.2-fold change compared to untreated are depicted. B) Results of Gene set enrichment in Gene Ontology Pathways. Differential gene analysis was performed using R package limma(2) with a p-value 0.05 adjusted with Benjamini-Hochberg multiple testing correction approach.

Combined administration of patient derived rFSGS/CD40autoAb and suPAR resulted in 1,783 gene expression changes. The gene enrichment showed regulation of megakaryocyte differentiation consistent with suPAR treatment. However, the addition of rFSGS/CD40autoAb did result in additional transcriptional changes (*FAP, VIT, ITGA7* among others) leading to pathways associated with cell adhesion (**Figure 3**). The largest set of transcriptionally altered genes was observed in this TNF treated group (5,094 genes) presumably due a more widespread effect of TNF treatment, but many of these pathways were also observed with rFSGS/CD40autoAb and suPAR treatment. The significant pathways associated with TNF treatment were broadly associated with regulation of the immune system. In addition, TNF also induced the expression of *APOC1, APOE, PLA2G2A* and *APOA1* associated with protein lipid complex remodeling (**Figure 3A and 3B**). A role for renal TNF in lipid dependent podocyte injury has been previously described [22].

### Identification of genes and pathways uniquely activated by CD40Ab and suPAR

As stated above, many of the genes that are transcriptionally altered in response to various stimuli are non-specific to FSGS. After conducting the differential gene expression, unique genes were selected using the Gene list Venn diagram software. There are 73 and 432 genes uniquely altered in response to rFSGS/CD40autoAb or suPAR alone respectively. Simultaneous treatment of podocytes with suPAR and rFSGS/CD40autoAb caused alterations in 511 unique genes. The largest subset of 3281 uniquely altered genes was associated with TNF stimulation (**Figure 4A**). 125 upregulated genes were common to all stimulations and associated with neutral lipid catabolic process, regulation of dendrite extension among others (**Figure 4B**). Interestingly, the 70 genes that are downregulated across all treatments did not lead to any known pathway enrichment. The top ten unique genes activated in response to each factor has been shown in the heatmap in **Figure 5**. Uniquely activated genes in response to combined treatment with suPAR and rFSGS/CD40autoAb included the TNF receptor superfamily member 25 (*TNFRSF25*) which is a well-known mediator of NFκB to regulate apoptosis [31].

**Figure 4.**
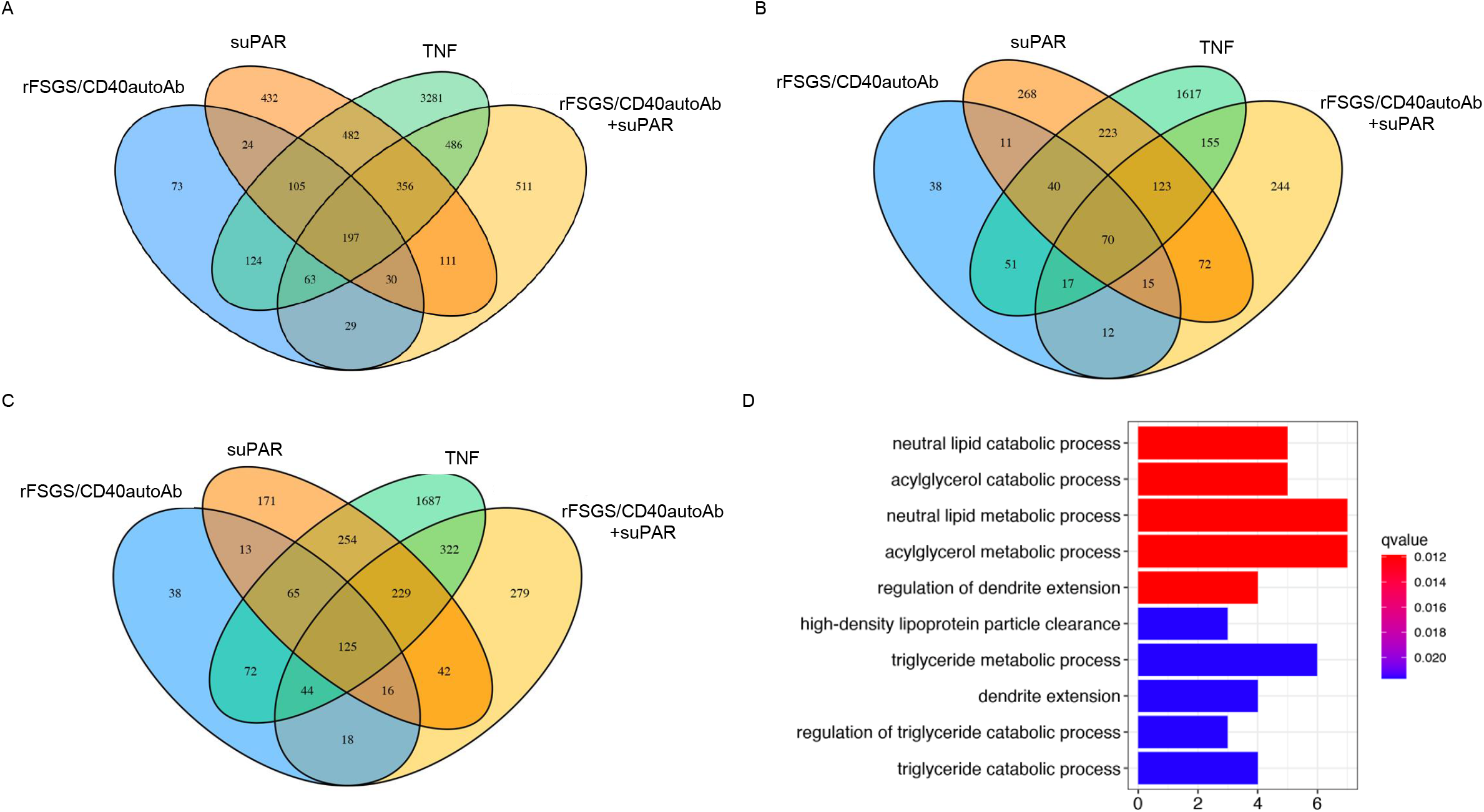
Characterization of genes and pathways uniquely activated by CD40Ab and suPAR. A) Venn diagram showing all, common and unique genes transcriptionally altered in podocytes in response to treatment with CD40autoAb purified from sera of FSGS patients with recurrence of FSGS after kidney transplant (rFSGS/CD40autoAb), suPAR, combination of rFSGS/CD40autoAb with suPAR or TNF alone as described in materials and methods. B) Profile of downregulated genes in podocytes after each treatment shows that 70 genes were downregulated across all treatments. However, gene enrichment analysis did not reveal any significant pathway enrichment. C) Upregulated genes in podocytes after aforementioned stimulations shows 125 common upregulated genes stimulated with CD40Ab, suPAR or CD40Ab and suPAR together. D) Pathways associated with genes upregulated in C. Gene sets were analyzed for enrichment in Gene Ontology biological pathways. Genes with more than 1.2 log fold change were included in the analysis and enriched pathways with q-value < 0.05 were accepted as significant.

**Figure 5.**
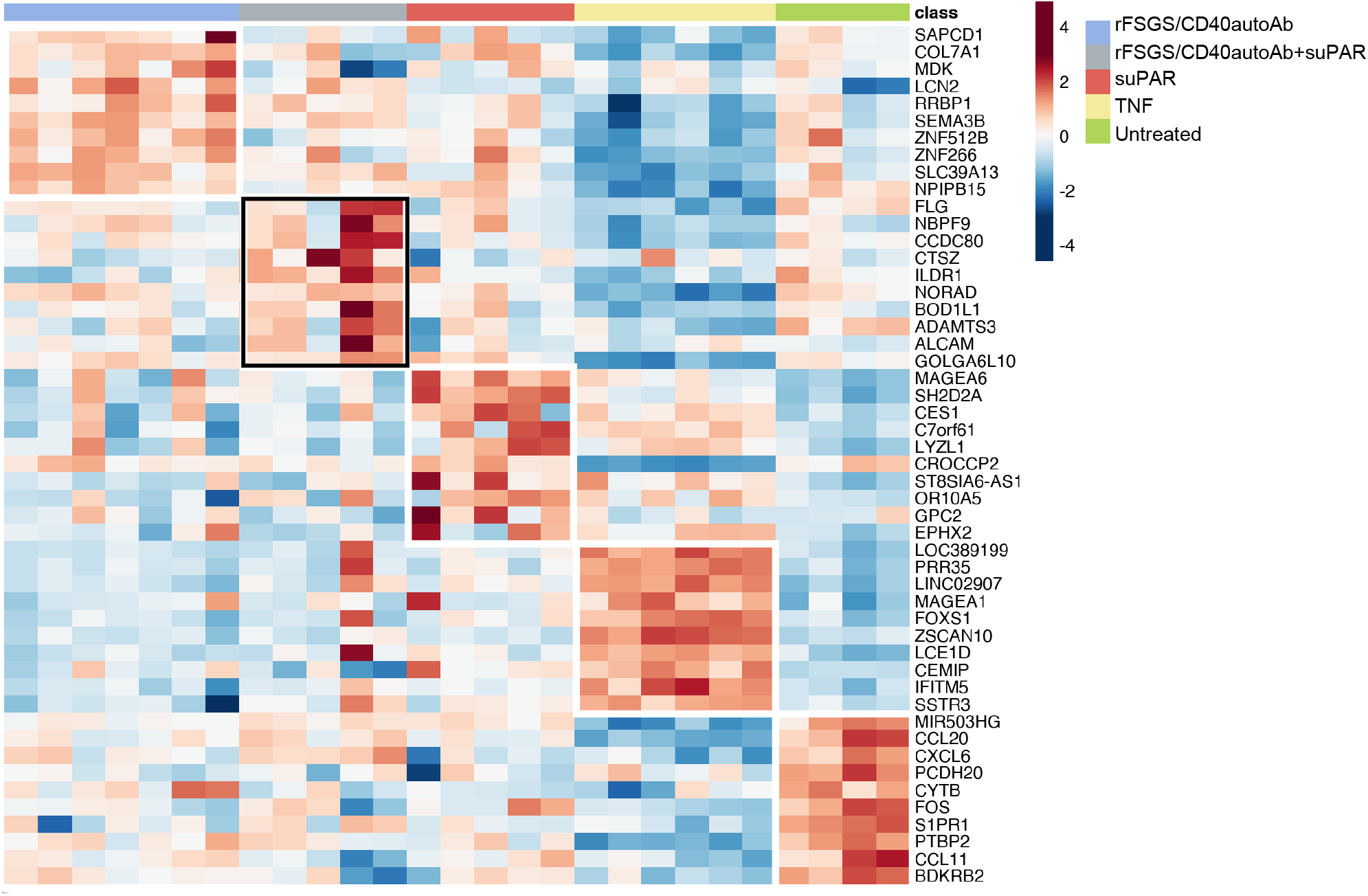
Identification of unique genes activated in response to various stimuli. To generate the stimulus-specific gene list, differential gene expression was conducted as described previously and unique genes were selected using the Gene list Venn diagram software.

### CD40 expression is downregulated in podocytes after treatment with suPAR and rFSGS-CD40Ab

Delville et al proposed a panel of seven antibodies (CD40, PTPRO, CGB5, FAS, P2RY11, SNRPB2 and APOL2) in the pre-transplant sera of FSGS patients that could predict posttransplant FSGS recurrence with 92% accuracy [7]. We tested if any of these genes were transcriptionally altered after podocyte injury in our study. Interestingly, *CD40* levels were downregulated after treatment with TNF, rFSGS/CD40autoAb or suPAR and could be rescued after treatment with commercial CD40 monoclonal antibody (**Figure 6**).

**Figure 6.**
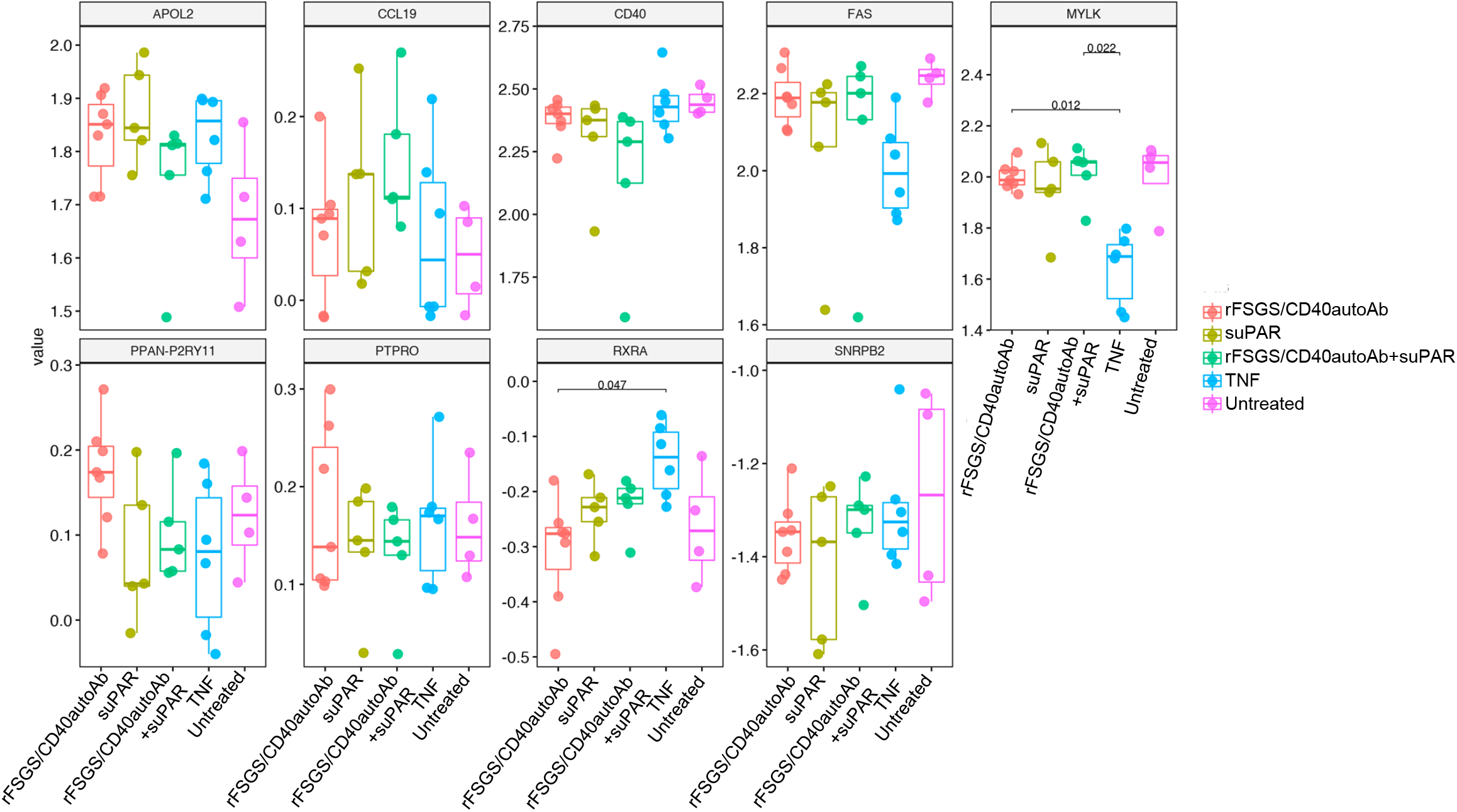
Transcriptomic regulation of antigens was previously described to be associated with recurrence of FSGS after kidney transplant (Delville et al). The expression level of indicated genes was compared in podocytes treated with CD40autoAb purified from sera of FSGS patients that recur after kidney transplant (rFSGS/CD40autoAb), suPAR, the combination of rFSGS/CD40autoAb and suPAR and TNF alone.

## Discussion

In this study, we aimed to further elucidate the role of CD40 and suPAR in FSGS and its recurrence through podocyte injury mechanisms. Here, we validate two novel antibodies against CD40 and suPAR as potential therapeutics for reversing podocyte injury associated with rFSGS. Furthermore, we identify molecules and pathways uniquely associated with rFSGS/CD40autoAb, or suPAR or their combination. The specific blockade of each of these pathways can be further evaluated for abrogating podocyte injury. Despite the morbidity of rFSGS, there are as yet, no specific therapies for the prevention or rapid reversal of podocyte injury after recurrence of FSGS after kidney transplantation. Therefore, an unmet need exists to develop and understand the mechanisms of novel therapeutic approaches to prevent or slow FSGS disease progression. suPAR and autoantibodies to several antigens have been proposed as biomarkers and possible therapeutic targets, but the heterogeneity of FSGS and the variability of patient specific responses to therapies have made substantive advances in therapeutic management extremely challenging.

Here, we attempt to understand podocyte injury in FSGS by the use of an established human podocyte *in vitro* model [7, 8], treated with human autoantibodies, suPAR and blocking target-specific antibodies. This model is relevant as the regulation of the podocyte actin cytoskeleton is crucial for the maintenance of glomerular filtration function and injury results in a shift towards increased motility of *in vitro* podocytes, reflected by foot process effacement *in vivo* [14]. This study supports that there is a pathogenic role for several circulating factors in rFSGS sera as contractile stress fibers in cultured podocytes are lost after treatment with whole sera and CD40autoAb isolated from rFSGS patients, but not with sera from FSGS patients that do not recur after kidney transplant. We assessed two human blocking antibodies-anti-uPAR antibody (2G10) and anti-CD40 antibody (BMS-986090), and both were found to prevent podocyte injury as mediated by rFSGS patient sera. Earlier studies have proposed that a full complex of uPA-uPAR binds to and activates β3-integrin on podocyte foot processes. This results in the activation of small GTPase Rac, induction of podocyte migratory phenotype and FSGS like phenotype. The anti-uPAR antibody, 2G10 blocks uPA-uPAR interaction suggesting that 2G10 mediated inhibition of podocytopathy progression involves an altered conformation of uPAR that cannot bind to and activate β3-integrin. At this time, the mechanism of rFSGS/CD40autoAb mediated podocyte injury as well as that of the antagonistic anti-CD40 antibody-mediated rescue of podocyte stress fiber loss induced by patient sera is not understood. The therapeutic effect of commercial monoclonal CD40 blocking antibody as opposed to the pathogenic effect of activating rFSGS/CD40autoAb isolated from patients has suggested that the two antibodies activate different signaling pathways and likely compete with CD40L for CD40 binding [7, 8]. In this study, different transcriptional changes in human podocytes to these CD40 antibodies can be observed. We have previously shown that higher concentrations of pre-transplant suPAR and rFSGS/CD40autoAb confer an increased risk of recurrence of FSGS after transplantation [7, 32]. We hypothesize that CD40autoAb synergizes with suPAR in causing podocyte injury by directly facilitating its binding with β3-integrin, the antagonistic anti-CD40 antibody might cause a conformational change in CD40 thus interfering with the synergy.

To further understand the molecular mechanism of podocyte injury induced by each of the aforementioned pathogenic factors and the pathways targeted by uPAR or CD40 blockade, we profiled the podocyte transcriptome after different stimulatory conditions. TNFα-treated podocytes served as a pre-established model of inflammatory injury and demonstrated dominant expression of multiple Apolipoprotein family members (*APOC1, APOE* and *APOA1*) [22]. This may be a non-specific final common pathway for podocyte injury from different triggers. Apolipoproteins have previously been reported in the context of FSGS as a modified form of Apolipoprotein A1 (ApoA-Ib) was detected in the urine of relapsing FSGS patients and later validated in an independent cohort as a potential biomarker to predict FSGS recurrence regardless of proteinuria status [33]. Individuals with genetic variants in the Apolipoprotein L1 gene (*APOL1*) have a greatly increased risk of hypertension-associated end-stage renal disease, focal segmental glomerulosclerosis (FSGS), and human immunodeficiency virus (HIV)-associated nephropathy [34–36]. Our earlier studies show that podocyte injury in response to combined suPAR and rFSGS/CD40autoAb is greater than the individual administration of each factor [7, 8]. Individual stimulation with rFSGS/CD40autoAb causes predominant activation of type I interferon signaling and genes involved in podocyte foot process effacement such as Nck2 [37]. However, suPAR alone primarily results in upregulation of genes involved in megakaryocyte differentiation. Common pathways perturbed across both suPAR and rFSGS/CD40Ab individual stimulations were associated with neutral lipid catabolic processes, cell adhesion mediated by integrins and included genes such as *LCN2* (Lipocalin 2). LCN2 has been proposed as a blood and urine biomarker for acute kidney injury (AKI) [38]. However, our data suggests that when combined with suPAR, rFSGS/CD40Ab mediates the induction of *TNFRSF25* (TNF receptor superfamily member 25) and several apolipoproteins, especially *APOL2. APOL2* forms a tripartite complex with αvβ3 complex in the presence of suPAR causing podocyte injury [39]. Therefore, it is likely that the formation of CD40autoAb and increased suPAR may be components of the two different hits which may be needed for the development or augmentation of renal injury mechanisms leading to FSGS.

Delville et al[7] proposed a panel of seven antibodies (CD40, PTPRO, CGB5, FAS, P2RY11, SNRPB2 and APOL2) in the pre-transplant sera of FSGS patients that could predict posttransplant FSGS recurrence with 92% accuracy. While there can be many reasons for the formation of autoantibodies, we tested if any of these genes were transcriptionally altered after podocyte injury in our study. We found that many of the apolipoproteins were dysregulated as described above but none of the other markers were transcriptionally altered apart from CD40. Interestingly, CD40 levels were downregulated after treatment with TNF, rFSGS/CD40autoAb or suPAR and could be rescued after treatment with commercial CD40 monoclonal antibody. This suggests that CD40-CD40L signaling is required for the maintenance of podocyte structure function and pathogenic factors act by abrogating this signaling albeit through different mechanisms.

In summary, through this study, we have been able to make several interesting observations related to the role of suPAR and CD40 and their antibodies in the recurrence of FSGS. By utilizing a human in vitro podocyte injury model, we examined the impact of two pathogenic factors (CD40 autoantibodies and suPAR) and their respective blockade antibodies on podocyte injury and alterations in podocyte mRNA levels. We can observe the synergy between rFSGS/CD40autoAb and suPAR in the development of podocyte injury and pathways that maybe of relevance to that injury in rFSGS. Additional *in vivo* studies focusing on suPAR, CD40autoAb and other pathogenic factors as well as new target-specific blocking antibodies for therapeutics will help further elucidate and unravel the different and myriad mechanisms of pathogenicity of rFSGS after kidney transplantation.

## Acknowledgments

The work presented in this manuscript was supported by funds from the NIDDK/NIH DK DK109720 to MS. The authors acknowledge help from Mr. Dane Munar for manuscript submission.

## Footnote

- **Reporting checklist**
- **Data sharing statement.** Microarray data used in this study will be made publicly available through NIH GEO repository
- **Conflict of Interest**: The other authors declare that the research was conducted in the absence of any commercial or financial relationships that could be construed as a potential conflict of interest.
- **Ethical Statement.** The authors are accountable for all aspects of the work in ensuring that questions related to the accuracy or integrity of any part of the work are appropriately investigated and resolved.

---

**Table S1.** Demographic information for patients with biopsy-confirmed recurrence of FSGS after transplant included in the study.

**Figure S1.** Treatment with sera from different patients with recurrence of FSGS (rFSGS) causes injury in podocytes which is not affected by treatment with a control human IgG. A) A representative image showing intact stress fibers in podocytes after treatment with sera from control patients (end-stage renal disease due to non-FSGS causes). However, sera from rFSGS patients lead to stress fiber loss which cannot be rescued by a control human IgG. B) Quantification of stress fibers positive cells show a significant reduction in a number of stress fiber positive podocytes after treatment with sera from rFSGS patients compared to that in control sera-treated podocytes and remains unchanged after pretreatment with control IgG.

**Figure S2.** Patient-derived CD40Ab causes injury to podocytes via a CD40-mediated pathway as it can be blocked by a commercial monoclonal anti-CD40 antibody (CD40mAb). Perturbations in podocyte transcriptome and biological pathways induced by FSGS associated circulating factors

