## Supplemental Table S1, Figure S1, and Figure S2 for "Perturbations in podocyte transcriptome and biological pathways induced by FSGS associated circulating factors"

### Supplementary materials

Table S1. Demographic information for patients with biopsy confirmed recurrence of FSGS after transplant included in the study.

| | Patient 1 | Patient 2 | Patient 3 |
|---|---|---|---|
| Age | 31 | 21 | 51 |
| Gender | M | F | M |
| Ethnicity | Caucasian | Caucasian | Asian |
| Living/deceased donor | Deceased | Living | Living |
| No. of days before recurrence of FSGS post Tx | Immediately | 7 | 195 |
| Maintenance Immunosuppression | | Simulect, MMF/MPA | Simulect, TAC/MMF |
| Maximum protein/creatinine at the time of recurrence | | 11,078 | 375 |
| Lowest Serum Albumin at the time of recurrence | | 5.44 | 0.91 |

### Figure S1

A

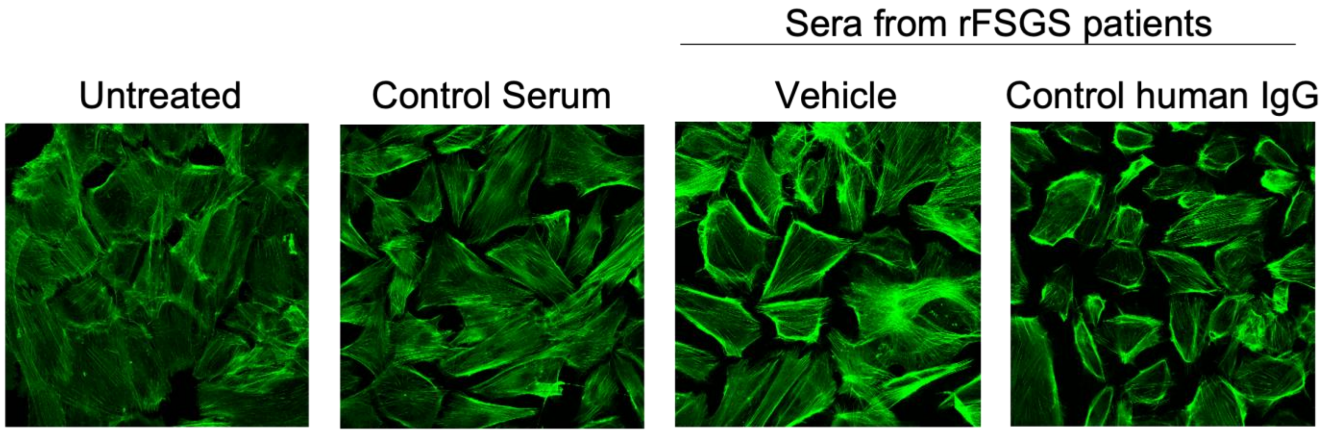

Sera from rFSGS patients
Untreated
Control Serum
Vehicle
Control human IgG


B

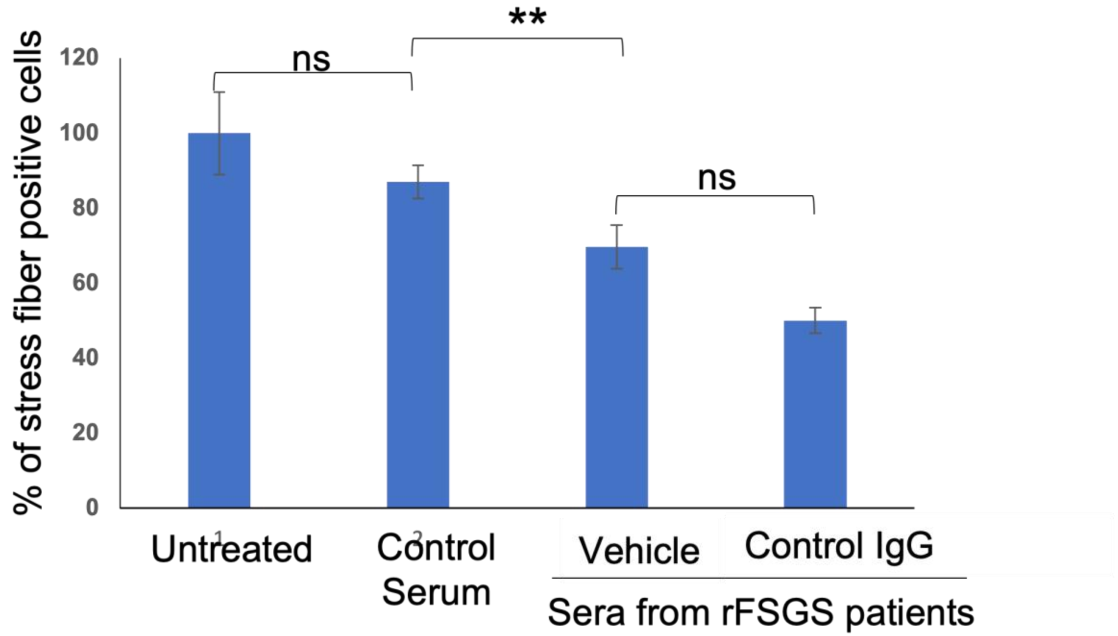

% of stress fiber positive cells
0
20
40
60
80
100
120
ns
**
ns
Untreated
Control Serum
Vehicle
Control IgG
Sera from rFSGS patients

### Figure S2

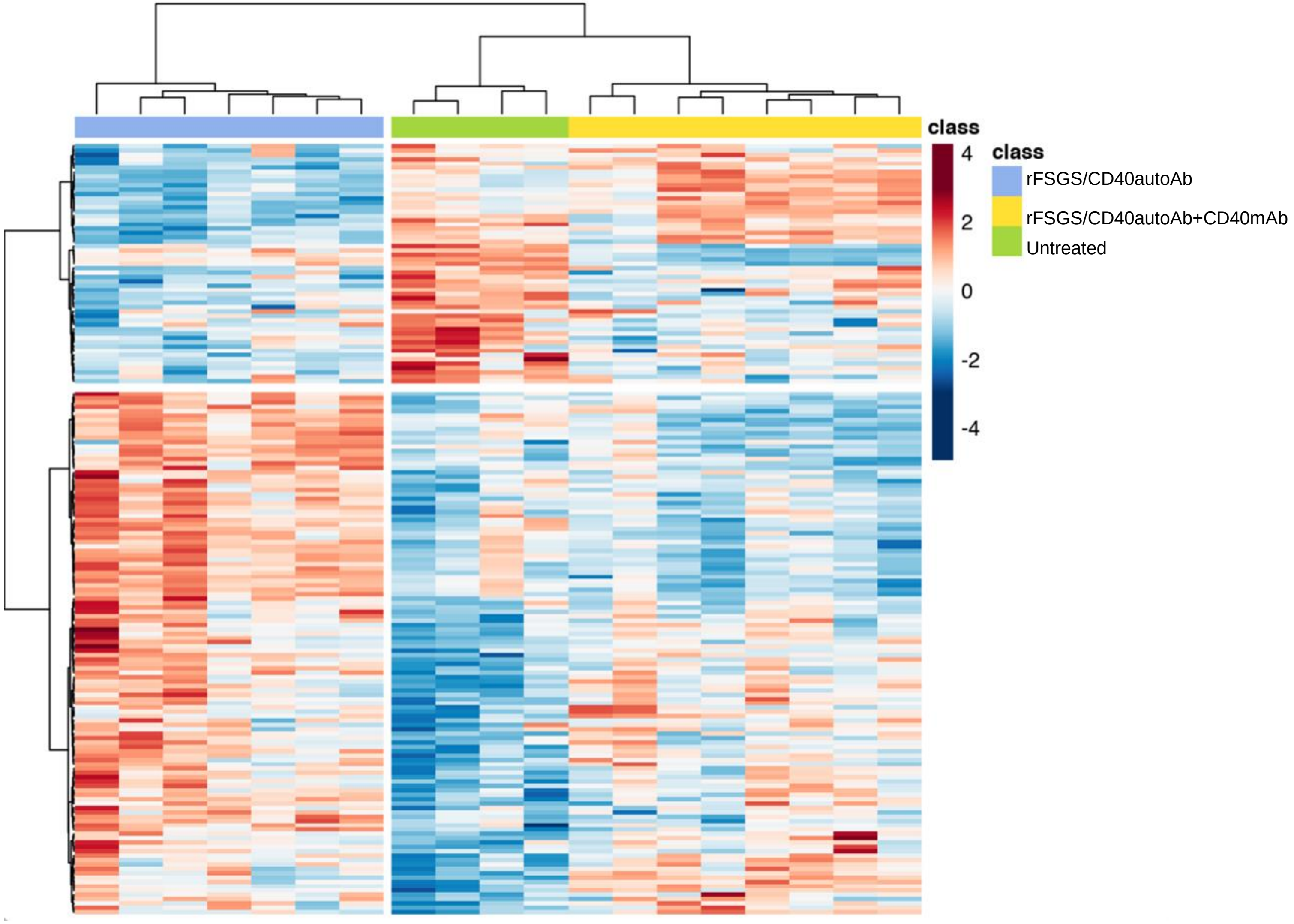

class
4
2
0
-2
-4
class
rFSGS/CD40autoAb
rFSGS/CD40autoAb+CD40mAb
Untreated
